# A Manually Curated Metabolic Model of *Alteromonas macleodii* MIT1002 and its Ecological Implications

**DOI:** 10.64898/2026.11.16.767100

**Authors:** Helen Scott, Devlin Moyer, Osnat Weissberg, Franziska Kratzl, Waseem Bashir Valiya Kalladi, Christa B. Smith, Rogier Braakman, Mary Ann Moran, Daniel Sher, Daniel Segrè

## Abstract

Heterotrophic bacteria are a vital part of the marine carbon cycle. They interact with phytoplankton and rapidly process fixed carbon, respiring it back to CO_2_. Among heterotrophs, *Alteromonas macleodii* is notable for its colonization of particulate material and its high growth and carbon utilization rates and is commonly used as a model system to study metabolic interactions with the abundant cyanobacterium *Prochlorococcus*. Understanding how these traits emerge from the interplay of hundreds of reactions requires a formal encoding of the organism’s metabolic network. Here, we present a metabolic model of *A. macleodii* MIT1002, built in a traceable and systematic way to recreate experimental growth phenotypes on 51 unique media compositions. We used this model to generate testable hypotheses about the role of sodium in the electron transport chain, the catabolic relevance of different glycolytic pathways, nitrogen recycling during cross-feeding with *Prochlorococcus*, and microbial carbon use efficiency.

**Importance:** The ocean is the largest carbon sink on the planet, but of the carbon that enters the ocean from the atmosphere, very little is stored over long periods of time. The majority is rapidly processed by heterotrophic bacteria that release CO_2_, thereby limiting the amount of carbon that can be stored. The rate at which heterotrophic bacteria take up organic carbon and release CO_2_ is determined, in part, by their metabolic networks. Despite the importance of heterotrophic metabolism, a detailed understanding of the metabolic network for a marine heterotrophic bacterium is rare. Here we created a detailed metabolic model for the bacterium *Alteromonas macleodii* MIT1002. We constructed this model in a documented and reproducible way and used the model to explore the role of MIT1002’s metabolism in carbon remineralization and interactions with the globally important cyanobacterium *Prochlorococcus*.

## Introduction

Heterotrophic bacteria play a key role in the marine carbon cycle through the consumption, transformation, and remineralization of organic carbon (Fig. 1A). In the surface ocean, heterotrophic bacteria rapidly consume the labile dissolved organic carbon (DOC) produced by phototrophs and utilize it for growth and energy, releasing much of it back to CO_2_ (1, 2). This high turnover rate limits the total amount of carbon that can be stored long-term in the ocean. In addition to their direct participation in the marine carbon cycle through remineralization (1–3), surface-water heterotrophs can also affect the rate of primary production through the regeneration of inorganic nutrients (4) and their interactions with phototrophs, which can be both positive (e.g., supplying vitamin B12 (5)) and negative (e.g., producing algicides (6)).

**Figure 1:**
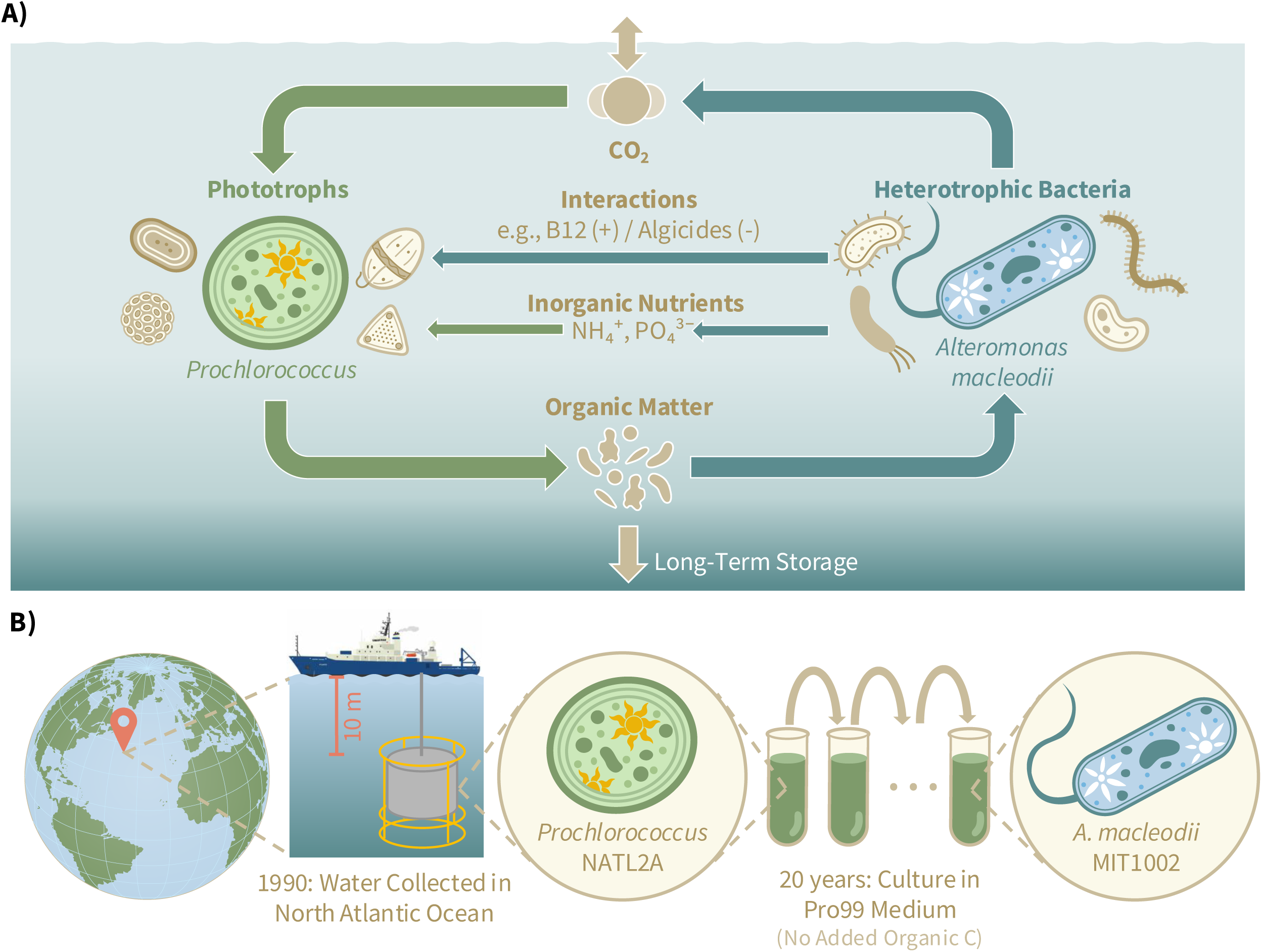
A) The role of heterotrophic bacteria in the marine carbon cycle. CO_2_ is fixed into organic carbon through photosynthesis by phototrophic microorganisms (e.g., *Prochlorococcus*). Organic matter produced by phototrophs is selectively taken up by heterotrophic bacteria (e.g., *Alteromonas macleodii*) and remineralized into CO_2_. The fast uptake and remineralization of organic carbon by heterotrophic bacteria limits the total amount of carbon that can be stored long-term in the ocean. Heterotrophs also affect the rate of primary production through the regeneration of inorganic nutrients (e.g., NH₄⁺, PO₄³⁻), and through interactions with phototrophs, which can be positive (e.g., the production of vitamin B12) or negative (e.g., the production of algicides). B) A water sample was collected from the North Atlantic Ocean (approximate location shown on the globe) at a depth of 10 meters. *Prochlorococcus* NATL2A was isolated from this water sample and then maintained in a continuous enrichment culture in Pro99 medium. After 20 years, MIT1002 was isolated from this enrichment culture.

One heterotrophic bacterium notable for high carbon processing rates and interactions with phototrophs is *Alteromonas macleodii* (7). This species is copiotrophic, achieving high growth rates when nutrients are readily available, such as during a phytoplankton bloom (8) or on particulate organic matter (9). This high growth rate is due, in part, to the bacterium’s ability to utilize a wide variety of organic compounds as substrates (7, 10–12). *A. macleodii* is a widely used model system for studying interactions between heterotrophic bacteria and the globally important cyanobacterium *Prochlorococcus* (12–22). While several orders of magnitude less abundant than *Prochlorococcus* in the wild, *Alteromonas* is consistently enriched in *Prochlorococcus* laboratory cultures (23), reflecting their metabolic compatibility. Indeed, in controlled co-cultures, the two bacteria form a mutualistic relationship: *Prochlorococcus* provides *Alteromonas* with the necessary organic carbon to support growth, and *Alteromonas* promotes the survival of *Prochlorococcus* during periods of oxidative stress (21, 22), extended darkness (17, 20), and nitrogen starvation (16, 19).

There is a high degree of strain-level variability within *A. macleodii* due to its extensive flexible genome, which is concentrated in genomic islands that evolve through the exchange of mobile genetic elements (12, 24, 25). This genetic variability results in substantial metabolic diversity (26–28): different strains utilize different substrates (12, 29), and have varied interactions with other microbes, including *Prochlorococcus* (15, 19, 30–32). The focus of this study is *A. macleodii* strain MIT1002 (hereafter MIT1002), which was initially isolated from a *Prochlorococcus* (strain NATL2A) enrichment culture (33) (Fig. 1B).

The molecular-scale processes that drive MIT1002’s remineralization of organic carbon and its interactions with *Prochlorococcus* can be studied using genome-scale metabolic modeling, which translates environmental constraints and genome knowledge into estimates of metabolic fluxes (34–36). Genome-scale metabolic modeling consists of two phases: (i) the reconstruction of an organism’s metabolic network, also known as a genome-scale metabolic model (GEM), and (ii) the prediction of fluxes through that network, based on simplifying assumptions, such as steady state and optimality, often using Flux Balance Analysis (FBA; see Methods) (34–37). Through predictions of uptake and secretion rates, FBA can be used to generate testable hypotheses about important ecological processes, including the dependence of Carbon Use Efficiency (CUE) on organismal metabolism and environmental constraints (38), and the rise of metabolite-mediated cross-feeding between microbes (39, 40). The predictive ability of FBA is strongly influenced by the quality of the GEM (37, 41). In parallel to the improvement of algorithms for the automatic construction and refinement of GEMs (42–48) and the development of approaches to filter models by their consistency and repeatability (49, 50), manual curation is still widely considered to be the gold-standard for maximizing the predictive potential of a GEM (41, 51). However, manual curation is rarely pursued due to the significant time, labor, and data requirements of this approach.

We present here MIT1002-GEM, a metabolic model for *A. macleodii* strain MIT1002 which we refined using a traceable and verifiable curation protocol. Through curation of individual reactions and pathways, our model accurately recapitulates 51 out of 62 experimental growth phenotypes. Drawing from established best practices in software development (52), we tracked changes to the model using version control (i.e., Git) and automatically tested the model throughout the curation process (52–54). We also documented all curation decisions in a standardized and open manner. Manual curation of this model identified reactions not yet represented in biochemistry databases that would be missed using automated curation tools and which relate to key features of life in the ocean. We used the resulting model to study metabolic processes relevant to the ecological role of MIT1002, including the possible role of seemingly redundant glycolytic pathways, the utilization of sodium ions in the electron transport chain, and the specific nitrogen-recycling role *Alteromonas* may fulfill in its interaction with *Prochlorococcus*.

## Methods

### Genome-Scale Metabolic Model (GEM) Construction and Curation

#### Draft Model Construction with KBase and ModelSEED

We constructed the draft model from the MIT1002 RefSeq genome (NZ_JXRW01000001) using KBase (55) and the underlying ModelSEED biochemistry database (46, 56). We first re-annotated the genome using the RASTtk app “Annotate Microbial Genome with RASTtk – v.1.073”. Options selected were: annotate proteins kmer v2, annotate proteins Kmer v1, and annotate proteins similarity. We then used the RAST-annotated genome to build a model with the ModelSEED2 app, “MS2 – Build Prokaryotic Metabolic Models with OMEGGA (v2.0.0)”, with no gap-filling. The Gram-negative template for reconstruction was selected automatically by the app. The resulting model had 942 genes, 1,335 reactions, and did not grow on the KBase “Complete” medium. Our KBase narrative is publicly available at https://narrative.kbase.us/narrative/191918. After initial construction of the draft model, we downloaded the model from KBase as an XML file (SBML format) and conducted all further curation manually.

#### Version Control and Testing with GitHub

We saved the resulting SBML model file in a public GitHub repository (https://github.com/C-CoMP-STC/MIT1002-GEM/) to track all changes to the model (54, 57) and the specific version of the model used here, v4.0.2, is archived on Zenodo (58). Curators proposed changes to the model and documented their reasoning in a GitHub “pull request”. Upon each pull request, GitHub Actions automatically ran a series of tests and reports to check how the proposed changes affected the model (e.g., the model’s ability to match experimental growth phenotypes, the presence of erroneous energy generating cycles, mass/charge imbalanced reactions, etc.) and help curators identify subsequent targets for manual curation. For full documentation of the automated testing workflow see the code repository (58).

#### Manual Curation of the Biomass Reaction

Prior to gap filling, we curated the default gram-negative biomass function, with a specific focus on removing biomass components that are known not to be present in MIT1002. We removed 2-Demethylmenaquinone 8 (cpd15352_c0) and Menaquinone 8 (cpd15500_c0), as functional annotation of the MIT1002 genome performed with eggNOG (59) did not identify any menaquinone biosynthesis genes in either the canonical (menA-F and menH) or alternative (mqnA-D) pathways, and other *Alteromonas* species have been reported to rely on ubiquinone-8 as the main or sole respiratory quinone (60–62). We also removed Stearoylcardiolipin *(B. subtilis*) (cpd15793_c0) from the biomass reaction as it is a generic template, not specific to MIT1002. Fatty acids in *Alteromonas* species are dominated by fatty acids with 16-carbon chains, or non-saturated 18-carbon chains, rather than the saturated 18-carbon chain in stearoylcardiolipin (60–62). And finally, we removed vitamin B12 (cpd00166_c0) as MIT1002 is known to not produce it (63, 64). We also removed all biomass components that did not represent ordinary metabolites (e.g., ACP and peptidoglycan) as they were not producible by the model and relied on loops in the biomass reaction (see Fig. S1). We then rescaled the coefficients of the biomass function so that 1 mol of biomass was equivalent to 1 gDW of biomass. See Table S1 for the final biomass composition.

#### Manual Gap Filling

Gap filling was informed by 62 binary experimental growth phenotypes (Table 2). For each of the 62 conditions, we ran FBA once for each biomass component while maximizing its production to determine if that component was producible in that medium condition. If a biomass component was not producible, we investigated its biosynthetic pathway for gaps (i.e., missing reactions). After a gap was identified, we consulted relevant literature about that biosynthetic pathway to determine which enzyme(s) were likely responsible for catalyzing the missing reaction(s), then searched for genomic evidence for those enzymes in functional annotations of the MIT1002 genome performed by RefSeq (65, 66) and eggNOG (59). If no genes in MIT1002 were confidently annotated with relevant catalytic activities, we looked for reciprocal best BLAST hits between the MIT1002 genome and a model organism or a close relative of MIT1002’s where the latter gene was annotated with a relevant catalytic activity. If we could not find a relevant gene but had experimental evidence supporting the existence of that reaction in *A. macleodii* we added the reaction to the model with no associated gene.

#### Removing Thermodynamically Infeasible Loops

Additional curation was needed to ensure that no thermodynamically infeasible loops could be used for energy generation (67). We systematically detected loops in the model using MACAW (68) and analyzed them in detail by identifying and editing reactions that carried a very large flux relative to nutrient upper bounds (e.g., > 100 mmol/gDW/h). We removed problematic loops by either removing reactions entirely or imposing irreversibility. Constraints on thermodynamic reaction reversibility were informed by Gibbs free energy changes (ΔG) available in the ModelSEED database or through literature-derived empirical evidence. For a detailed example of a loop that was removed, see Fig. S2.

#### Adding Genes/Reactions from Pangenome Analysis

A previous study constructed an *Alteromonas* pangenome containing 339 genomes, including MIT1002, and inferred the presence of genes encoding enzymes capable of catalyzing particular metabolic reactions across all included genomes (69). Because the pangenome used a different genome assembly for MIT1002 (JGI Taxon ID: 2738541267) from the one used to construct the draft model, to compare the functional annotations of the genes inferred to be present in MIT1002 in this pangenome to the annotations in the MIT1002 genome used to create the draft model we used PSI-BLAST (BLAST+ 2.2.2.0) to map genes between the two genomes. Since the draft model was created with ModelSEED, but the pangenome analysis annotated genes with KEGG reaction IDs, we also translated KEGG reaction IDs to ModelSEED reactions, using the ModelSEED ID with the lowest number and a “status” of “OK” that was not annotated as obsolete in ModelSEED. There were 349 reactions inferred to be present in MIT1002 in the pangenome analysis. Of those reactions, 151 were already included in the draft model, and 82 were either redundant with reaction(s) already in the model or represented non-metabolic reactions (e.g. post-translational modifications, DNA/RNA modifications). We added the remaining 116 reactions to the model. The pangenome analysis associated some of the 151 reactions that were already present in the draft model with genes that had no specific annotated function in the MIT1002 genome used to make the draft model, so we expanded the gene-protein reaction rules for some of those 151 reactions based on the pangenome analysis.

### Flux Balance Analysis

Flux Balance Analysis (FBA), described in detail elsewhere (34, 70, 71), uses linear programming to predict a set of steady-state fluxes through a metabolic network (including uptake, secretion, and growth rate) that optimizes the flux through specific reactions in the network, given constraints on the fluxes through particular reactions (e.g. nutrient availability or reversibility). In FBA, the metabolic network from a GEM is encoded into a Stoichiometric Matrix (*S*), where columns represent reactions, and rows represent metabolites. Each entry in the matrix is the corresponding stoichiometric coefficient for that particular metabolite in that reaction. A vector of fluxes *v* (one flux for each reaction in the network) that satisfies the equation *Sv*=*0* is a set of steady-state fluxes through that network, where the sum of all fluxes consuming each metabolite is equal to the sum of all fluxes producing it. Each reaction’s flux is itself constrained by an upper (*v_i_*_, *max*_) and lower (*v_i_*_, *min*_) bound. The objective function can be any linear combination of fluxes (*c^T^v*), with each flux (*v_i_*) weighted by a coefficient (*c*_i_). Taken together, the standard form of FBA is:

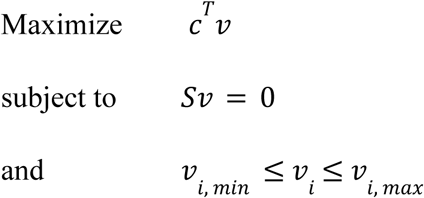

Parsimonious FBA (pFBA) is a commonly used extension to FBA that has been shown to more accurately predict fluxes (72). In pFBA, a secondary optimization minimizes the total sum of fluxes after fixing the objective (e.g., growth rate) at the optimal value.

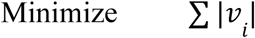

This additional optimization is advantageous because it reduces the number of alternative optimal solutions and is biologically grounded as it assumes that cells have evolved to minimize protein costs (73).

For this study, we ran all FBA and pFBA simulations in Python using COBRApy version 0.29.1 (74).

### Flux Visualization with Escher

To explore and visualize the fluxes predicted by FBA, we used Escher maps (75, 76). We constructed an Escher map of central metabolism for MIT1002. All Escher maps are saved as JSON files and are available as part of the model repository (58).

### Defining and Calculating CUE

CUE is defined as the fraction of the total amount of carbon taken up that remains within the system (77). Consistent with (77), we used the following definition:

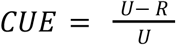

Where “R” is the flux of CO_2_ being released by the cell and “U” is the flux of all organic carbon taken up by the cell. These fluxes can be extracted from an FBA solution by multiplying the relevant exchange reaction fluxes by the number of carbon atoms in the exchanged metabolite (38). While this contrasts with typical definitions of growth yield it is the most relevant definition when studying how microbial ecosystems retain or respire carbon.

### Experimental Growth Phenotyping

In addition to compiling previously published experimental growth phenotypes for MIT1002 (12, 78, 79), we performed our own phenotyping experiments. We cultured MIT1002 for approximately 4 hours on rich medium (½ YTSS; yeast extract, tryptone, and artificial seawater) to early-exponential phase. Cells were then pelleted and washed in the minimal medium three times to remove any excess organic carbon. We used the defined Minimal Basal Medium (see Supplementary Material 1), amended with individual carbon sources to a concentration equivalent to 6, 8, or 12 mM carbon. Washed cells were added at 1% volume to growth medium in either 48- or 96-well plates and were grown at 28°C with constant shaking for up to 72 hours. The optical density at 600 nm (OD600) was measured on a Synergy H1 plate reader. For OD600 measurements for all substrates, see Supplementary Material 2 and Fig. S4.

## Results

We constructed and manually curated a genome-scale metabolic model (GEM) for *A. macleodii*MIT1002. As summarized in Table 1, the resulting model, MIT1002-GEM, has 1,426 reactions, 1,238 metabolites, and 949 genes, accounting for 23.7% of the 4,011 protein-coding genes (as annotated by RefSeq), and 89.4% of the 1,062 genes assigned an EC number by eggNOG-mapper v2.1.12. MIT1002-GEM accurately predicts if MIT1002 is capable of growing or not for 51 of the 62 experimentally tested media. Table 2 shows each of these 62 binary growth phenotypes, with the FBA-predicted growth rate on each.

**Table 1:** Statistics on model size and performance for the initial draft model reconstruction (v1.0.0) and the current, manually curated model (v4.0.2). The initial draft model reconstruction was made on KBase using the ModelSEED2 app, “MS2 – Build Prokaryotic Metabolic Models with OMEGGA (v2.0.0)”. Model performance was measured by the model’s ability to match experimentally determined growth phenotypes for 62 unique media compositions. The model is considered to “match” the experimental growth phenotype if it either predicts growth when growth was experimentally observed (true positive) or predicts no growth when no growth was observed (true negative).

|  | **Draft (v1.0.0)** | **Current (v4.0.2)** |
|---|---|---|
| **Model size** |  |  |
| Reactions | 1335 | 1426 |
| Metabolites | 1212 | 1238 |
| Genes | 942 | 949 |
| **Growth phenotypes (n = 62)** |  |  |
| Matches | 23 (37%) | 51 (82%) |
| Growth observed, growth predicted | 0 | 31 |
| Growth observed, no growth predicted | 39 | 8 |
| No growth observed, growth predicted | 0 | 3 |
| No growth observed, no growth predicted | 23 | 20 |

**Table 2:** Experimentally determined growth phenotypes and the FBA-predicted growth rate on the defined medium. Rows colored pink rows are mismatches (i.e., incorrectly predicting growth or no growth), and rows colored in gray are experimental phenotypes that are excluded from comparison (e.g., due to lack of a control). For each medium, the bound for carbon source uptake is set to 60 mmol C / gDW / h. The bound for each carbon source is set by normalizing that rate by the number of carbon atoms in the substrate being tested. All other media components (including nitrogen sources) are unlimited, and oxygen availability is limited to 20 mmol / gDW / h. Protons and water are also unlimited. Experimental growth phenotypes from (Halloran et al., 2026) were defined by testing if carrying capacity was significantly higher when A. macleodii MIT1002 cultures received the metabolite than when A. macleodii MIT1002 received no Prochlorococcus metabolite. For the FBA predictions, no alternative carbon sources were included in the ProMM medium. OD600 measurements for growth phenotypes from this study are available in Supplementary File S2.

| Base Medium | N Source(s) in Base Medium | Added Metabolite(s) | Experimental Growth | Reference | Reason for Exclusion | FBA Predicted Growth Rate | Result |
|---|---|---|---|---|---|---|---|
| Kratzl Base Medium |  | Ammonium, Succinate | No | (Kratzl, 2026) |  | 0.0 | True Negative |
| Kratzl Base Medium |  | Nitrate, Succinate | No | (Kratzl, 2026) |  | 0.0 | True Negative |
| Kratzl Base Medium |  | Urea, Succinate | No | (Kratzl, 2026) |  | 0.0 | True Negative |
| Kratzl Base Medium (with N-solution) | NH3, Nitrate | Alanine | Yes | (Kratzl, 2026) |  | 0.585 | True Positive |
| Kratzl Base Medium (with N-solution) | NH3, Nitrate | Alanine, Pyruvate | Yes | (Kratzl, 2026) |  | 0.766 | True Positive |
| Kratzl Base Medium (with N-solution) | NH3, Nitrate | Arginine | Yes | (Kratzl, 2026) |  | 0.613 | True Positive |
| Kratzl Base Medium (with N-solution) | NH3, Nitrate | Arginine, Pyruvate | Yes | (Kratzl, 2026) |  | 0.78 | True Positive |
| Kratzl Base Medium (with N-solution) | NH3, Nitrate | Cystine | No | (Kratzl, 2026) |  | 0.0 | True Negative |
| Kratzl Base Medium (with N-solution) | NH3, Nitrate | Cystine, Pyruvate | No | (Kratzl, 2026) |  | 0.565 | False Positive |
| Kratzl Base Medium (with N-solution) | NH3, Nitrate | Glutamine | Yes | (Kratzl, 2026) |  | 0.719 | True Positive |
| Kratzl Base Medium (with N-solution) | NH3, Nitrate | Glutamine, Pyruvate | Yes | (Kratzl, 2026) |  | 0.889 | True Positive |
| Kratzl Base Medium (with N-solution) | NH3, Nitrate | Lysine | Yes | (Kratzl, 2026) |  | 0.504 | True Positive |
| Kratzl Base Medium (with N-solution) | NH3, Nitrate | Lysine, Pyruvate | Yes | (Kratzl, 2026) |  | 0.692 | True Positive |
| Kratzl Base Medium (with N-solution) | NH3, Nitrate | Methionine | No | (Kratzl, 2026) |  | 0.0 | True Negative |
| Kratzl Base Medium (with N-solution) | NH3, Nitrate | Methionine, Pyruvate | Yes | (Kratzl, 2026) |  | 0.565 | True Positive |
| Kratzl Base Medium (with N-solution) | NH3, Nitrate | Proline, Pyruvate | Yes | (Kratzl, 2026) |  | 0.852 | True Positive |
| Minimal Basal Medium (Moran Lab) | NH3 | 3-hydroxybutyrate | Yes | This study |  | 0.55 | True Positive |
| Minimal Basal Medium (Moran Lab) | NH3 | Acetate | Yes | This study |  | 0.385 | True Positive |
| Minimal Basal Medium (Moran Lab) | NH3 | Acetoin | No | This study |  | 0.0 | True Negative |
| Minimal Basal Medium (Moran Lab) | NH3 | Alanine | Yes | This study |  | 0.585 | True Positive |
| Minimal Basal Medium (Moran Lab) | NH3 | Alginate | No | This study |  | 0.0 | True Negative |
| Minimal Basal Medium (Moran Lab) | NH3 | Allantoin | No | This study |  | 0.0 | True Negative |
| Minimal Basal Medium (Moran Lab) | NH3 | Asparagine | Yes | This study |  | 0.612 | True Positive |
| Minimal Basal Medium (Moran Lab) | NH3 | Choline | Yes | This study |  | 0.0 | False Negative |
| Minimal Basal Medium (Moran Lab) | NH3 | Citrate | No | This study |  | 0.0 | True Negative |
| Minimal Basal Medium (Moran Lab) | NH3 | Ectoine | Yes | This study |  | 0.0 | False Negative |
| Minimal Basal Medium (Moran Lab) | NH3 | Formate | No | This study |  | 0.0 | True Negative |
| Minimal Basal Medium (Moran Lab) | NH3 | Fructose | Yes | This study |  | 0.0 | False Negative |
| Minimal Basal Medium (Moran Lab) | NH3 | Fumarate | No | This study |  | 0.611 | False Positive |
| Minimal Basal Medium (Moran Lab) | NH3 | Galactose | Yes | This study |  | 0.73 | True Positive |
| Minimal Basal Medium (Moran Lab) | NH3 | Galacturonic Acid | Yes | This study |  | 0.645 | True Positive |
| Minimal Basal Medium (Moran Lab) | NH3 | Gluconate | Yes | This study |  | 0.0 | False Negative |
| Minimal Basal Medium (Moran Lab) | NH3 | Glucose | Yes | This study |  | 0.75 | True Positive |
| Minimal Basal Medium (Moran Lab) | NH3 | Glucuronic Acid | No | This study |  | 0.0 | True Negative |
| Minimal Basal Medium (Moran Lab) | NH3 | Glutamate | Yes | This study |  | 0.71 | True Positive |
| Minimal Basal Medium (Moran Lab) | NH3 | Glyceraldehyde 3-phosphate | No | This study |  | 0.0 | True Negative |
| Minimal Basal Medium (Moran Lab) | NH3 | Glycerol | Yes | This study |  | 0.766 | True Positive |
| Minimal Basal Medium (Moran Lab) | NH3 | Glycolate | No | This study |  | 0.0 | True Negative |
| Minimal Basal Medium (Moran Lab) | NH3 | Glyoxylate | No | This study |  | 0.0 | True Negative |
| Minimal Basal Medium (Moran Lab) | NH3 | Lactate | Yes | This study |  | 0.585 | True Positive |
| Minimal Basal Medium (Moran Lab) | NH3 | Leucine | Yes | This study |  | 0.499 | True Positive |
| Minimal Basal Medium (Moran Lab) | NH3 | Lysine | Yes | This study |  | 0.466 | True Positive |
| Minimal Basal Medium (Moran Lab) | NH3 | Mannuronic Acid | Yes | This study |  | 0.645 | True Positive |
| Minimal Basal Medium (Moran Lab) | NH3 | Phenylalanine | Yes | This study |  | 0.0 | False Negative |
| Minimal Basal Medium (Moran Lab) | NH3 | Proline | Yes | This study |  | 0.638 | True Positive |
| Minimal Basal Medium (Moran Lab) | NH3 | Putrescine | Yes | This study |  | 0.0 | False Negative |
| Minimal Basal Medium (Moran Lab) | NH3 | Pyruvate | Yes | This study |  | 0.565 | True Positive |
| Minimal Basal Medium (Moran Lab) | NH3 | Succinate | No | This study |  | 0.0 | True Negative |
| Minimal Basal Medium (Moran Lab) | NH3 | Sucrose | Yes | This study |  | 0.0 | False Negative |
| Minimal Basal Medium (Moran Lab) | NH3 | Valine | Yes | This study |  | 0.47 | True Positive |
| Minimal Basal Medium (Moran Lab) | NH3 | Xylose | No | This study |  | 0.674 | False Positive |
| ProMM | NH3 | 3-methyl-2-oxobutanoic acid | Yes | (Halloran, 2026) |  | 0.491 | True Positive |
| ProMM | NH3 | 3-methyl-2-oxopentanoic acid | No | (Halloran, 2026) |  | 0.0 | True Negative |
| ProMM | NH3 | 4-hydroxybenzoic acid | No | (Halloran, 2026) |  | 0.0 | True Negative |
| ProMM | NH3 | 4-methyl-2-oxopentanoic acid | No | (Halloran, 2026) |  | 0.0 | True Negative |
| ProMM | NH3 | Isoleucine | Yes | (Halloran, 2026) |  | 0.537 | True Positive |
| ProMM | NH3 | Kynurenine | No | (Halloran, 2026) |  | 0.0 | True Negative |
| ProMM | NH3 | Leucine | Yes | (Halloran, 2026) |  | 0.499 | True Positive |
| ProMM | NH3 | Thymidine | No | (Halloran, 2026) |  | 0.0 | True Negative |
| ProMM | NH3 | Valine | Yes | (Halloran, 2026) |  | 0.47 | True Positive |
| Seawater Medium | NH3 | Phenol | Yes | (Koch, 2020) |  | 0.265 | True Positive |
| Seawater Medium | NH3 | alpha-Mannan | Yes | (Koch, 2020) |  | 0.0 | False Negative |
| Minimal Basal Medium (Moran Lab) | NH3 | 6-Phosphogluconate | Unsure | This study |  | 0.778 | Unsure |
| Minimal Basal Medium (Moran Lab) | NH3 | DHPS (dihydroxypropanesulfonate) | Unsure | This study |  | 0.607 | Unsure |
| Minimal Basal Medium (Moran Lab) | NH3 | Glycerol-3-phosphate | Unsure | This study |  | 0.926 | Unsure |
| Minimal Basal Medium (Moran Lab) | NH3 | Glycine | Unsure | This study |  | 0.574 | Unsure |
| Minimal Basal Medium (Moran Lab) | NH3 | Phosphoenolpyruvate | Unsure | This study |  | 0.705 | Unsure |
| Minimal Basal Medium (Moran Lab) | NH3 | Tyrosine | Unsure | This study |  | 0.327 | Unsure |
| Kratzl Base Medium |  | Ammonium, Glucose | Yes | (Kratzl, 2026) | control_failed | 0.75 | Excluded |
| Kratzl Base Medium |  | Ammonium, Pyruvate | Yes | (Kratzl, 2026) | control_failed | 0.565 | Excluded |
| Kratzl Base Medium |  | Nitrate, Glucose | Yes | (Kratzl, 2026) | control_failed | 0.647 | Excluded |
| Kratzl Base Medium |  | Nitrate, Pyruvate | Yes | (Kratzl, 2026) | control_failed | 0.486 | Excluded |
| Kratzl Base Medium |  | Urea+Pyruvate | No | (Kratzl, 2026) | control_failed | 0.0 | Excluded |
| Kratzl Base Medium |  | Urea, Glucose | Yes | (Kratzl, 2026) | control_failed | 0.0 | Excluded |
| Kratzl Base Medium (with N-solution) | NH3, Nitrate | Proline | No | (Kratzl, 2026) | conflicting_reports | 0.687 | Excluded |

Achieving this level of accuracy required extensive manual curation, in a process spanning over one year and 100 GitHub pull requests. Throughout the manual curation process we evaluated the model’s performance by the number of experimental phenotypes it correctly predicted (Fig. 2A) and the number of reactions with unreasonably high fluxes in pFBA solutions (Fig. 2B), indicative of thermodynamically infeasible loops that could lead to an artificially large growth rate (67). Notably, model performance did not improve monotonically, with some changes to the model resulting in worse model performance (Fig. 2). The use of GitHub for version control was instrumental in detecting and correcting these detrimental changes.

**Figure 2:**
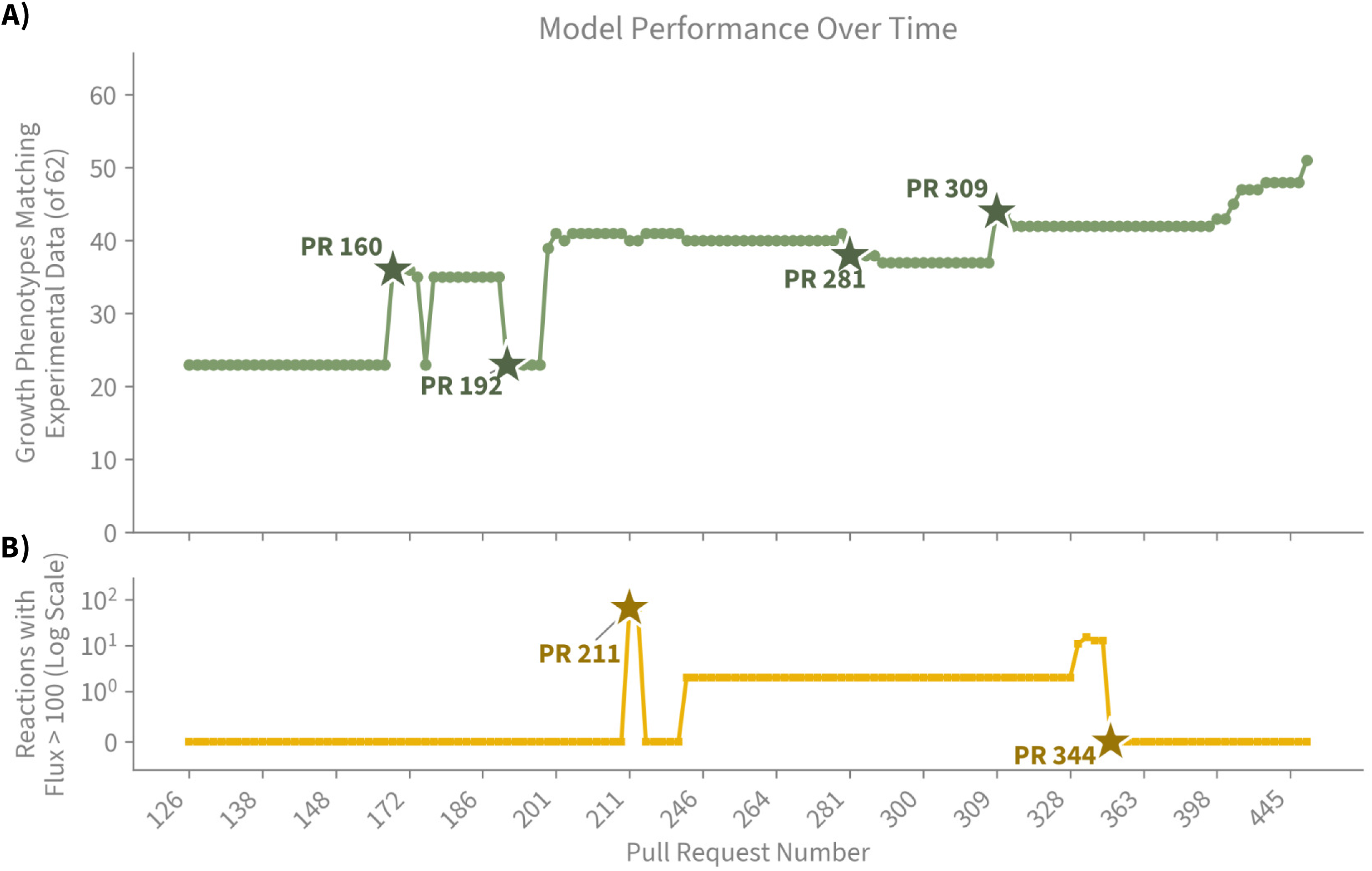
Model Performance over time from initial draft model construction to the current release presented here, as judged by A) the number of growth phenotypes matching experimental data, and B) by the number of unique reactions across all growth conditions that carry a flux greater than 100 mmol/gDW/h. Model performance over time shows a non-linear and non-monotonic trend over time. Pull Requests (PRs) that significantly changed model performance are marked with a star: PR 160) added two reactions to connect dead-end metabolites to glyoxylate, PR 192) removed duplicate thiamine reaction, PR 211) removed reactions with glucose anomers, PR 281) standardized amino acid transporters (no longer grows on leucine and isoleucine), PR 309), adde d Na^+^ symporters for leucine and isoleucine, PR 344) fixed the GAPDHN infeasible loop.

### Manual Curation of the Electron Transport Chain

During the manual curation process, we found a gap in the electron transport chain, with the model missing the reaction catalyzed by Complex I (NADH:ubiquinone oxidoreductase). No genes in the MIT1002 genome were associated with this reaction. However, functional annotation performed with eggNOG (59) annotated all 6 subunits of the multisubunit NQR complex (nqrA-nqrF) responsible for catalyzing the sodium-translocating NADH:quinone oxidoreductase (Na^+^-NQR). This is a known alternative to the proton-translocating reaction and is widespread among marine bacteria (80). Because this reaction is not present in the ModelSEED database, it had been left out of the draft model. We formulated a new reaction in MIT1002-GEM called “ec7211” with a stoichiometry consistent with (81):

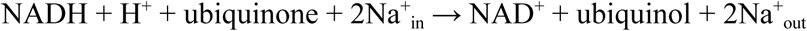

With the addition of the Na^+^-NQR reaction, the model was capable of generating and utilizing a sodium motive force (SMF), analogous to the proton motive force (PMF), for energy generation. The SMF has previously been described for marine bacteria (80–82), and its presence is tied to the marine environment. Because the pH of seawater is relatively high (∼8.0), a microbe’s cytosol is more acidic than the environment, making the generation and maintenance of a PMF difficult (80). However, high concentrations of sodium in seawater allow for the generation of a SMF (81), that can be used to drive ATP synthesis, transport, and motility.

Different reactions involved in generating and dissipating the SMF and PMF are shown in Figure 3A, and the fluxes through those reactions on different substrates are shown in Figure 3B-C. Figure 3B shows that the SMF is generated entirely by Na^+^-NQR and is primarily consumed by the Na^+^/H^+^ antiporter, effectively trading SMF for PMF. Growth on carbon sources that are imported into the cell with a sodium symporter (i.e., amino acids) also utilizes a fraction of the generated SMF for transport, reducing the flux through Na^+^/H^+^ antiporter, and therefore the amount of PMF generated (Fig. 3B).

**Figure 3:**
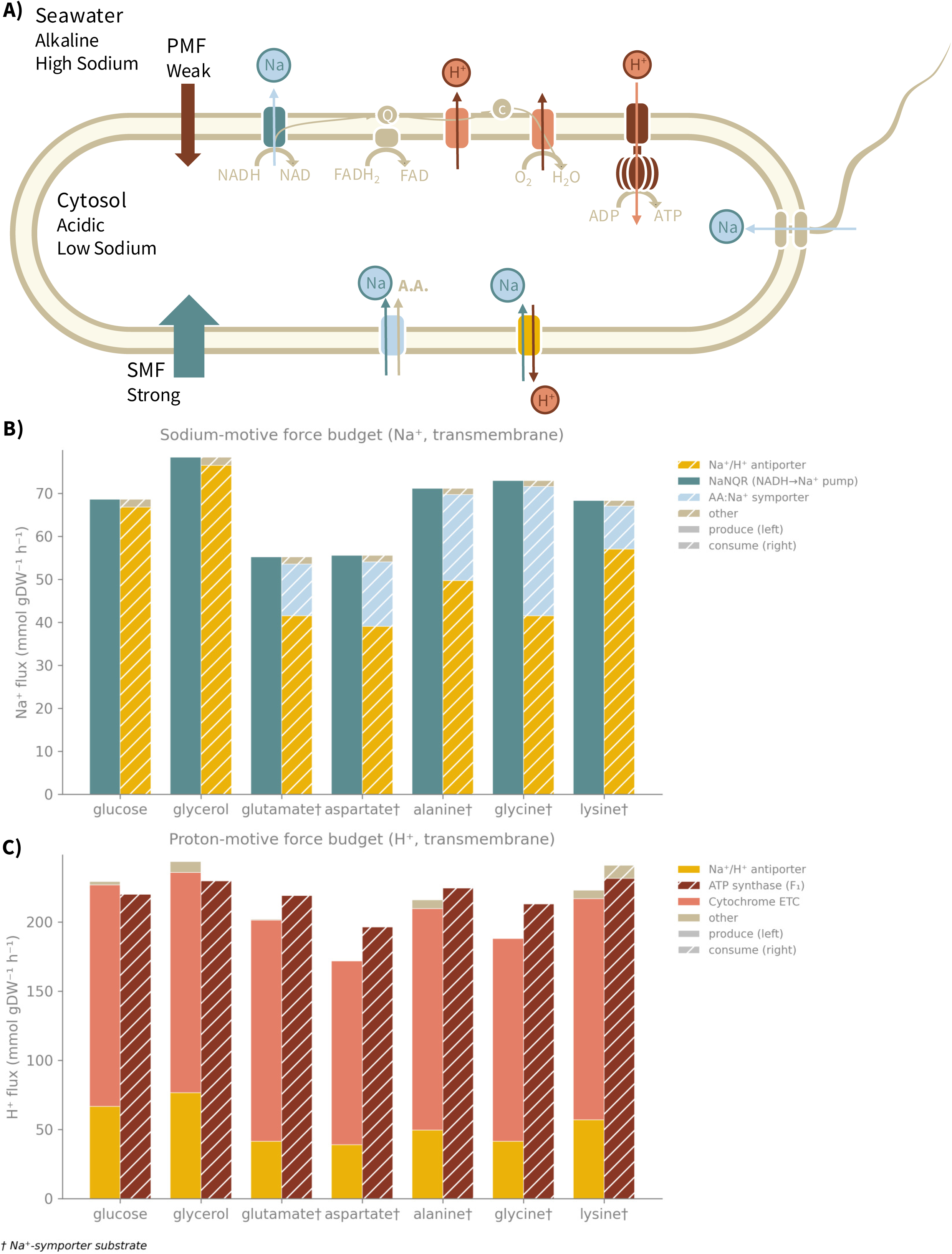
The Sodium Motive Force. A) A schematic diagram of key reactions responsible for generating and utilizing the sodium motive force (SMF) and proton motive force (PMF). The PMF is weak for marine bacteria due to the relative alkalinity of seawater. However, the high salinity of seawater allows for the maintenance of a strong SMF. The PMF and its reactions are shown in red/pink, and the SMF and its reactions are blue. The electron transport chain (ETC) is shown at the top of the bacterial diagram. The ETC begins with a Na^+^-translocating NADH:quinone oxidoreductase (Na^+^-NQR, shown in teal), which generates the SMF by pumping Na^+^ out of the cell. The remaining steps of the ETC pump H^+^s, as is typically described. ATP synthase exclusively uses the PMF by importing H^+^s into the cell. Other reactions that consume the SMF include amino acid symporters (blue reaction on the bottom), and the SMF also may power MIT1002’s flagellum. A Na^+^/H^+^ antiporter (shown in yellow) can be used to exchange SMF for PMF or vice versa. B) Sodium fluxes into and out of the cell when grown on different carbon sources. Each carbon source has stacked bars of the Na^+^ flux out of the cell, producing the SMF (left), and into the cell, consuming the SMF (right). For all substrates, the only reaction producing the SMF is the Na^+^-NQR. For glucose and glycerol, the majority of the SMF is consumed by the Na^+^/H^+^ antiporter. However, for growth on amino acids, some of the SMF is also consumed by transport. C) Proton fluxes into and out of the cell when grown on different carbon sources. Each carbon source has stacked bars of the H^+^ flux out of the cell, producing the PMF (left), and into the cell, consuming the PMF (right). For all substrates, the PMF is generated by the Na^+^/H^+^ antiporter and by the H^+^ pumping reactions of the ETC. For most carbon sources, the exclusive use of the PMF is ATP synthesis.

An additional possible role for the SMF is to power MIT1002’s flagellum. We believe MIT1002’s flagellum may be Na^+^-driven because it has been described as single and polar (28), consistent with other Na^+^-powered flagella (80), and RefSeq and eggNOG both annotated the MIT1002 genome with the PomA gene for the Na^+^-driven flagellum. While FBA does not typically model motility, it can incorporate maintenance costs. With the addition of a pseudoreaction that imports Na^+^ into the cell, as the flagellum would, the metabolic model predicts a tradeoff between energy generation and motility (Fig. S5).

### Ecological Implications of MIT1002 Metabolism

Once we were satisfied with the model’s performance, we used MIT1002-GEM to investigate metabolic traits of MIT1002. Among the hundreds of metabolic pathways one could analyze in detail, we chose a few examples motivated by their relevance to MIT1002’s role in the marine carbon cycle (highlighted in the metabolic map in Figure 4). In particular, given *A. macleodii*’s ability to grow on many different compounds, we used MIT1002-GEM to provide insight into how the presence of specific catabolic pathways may relate to the availability of environmental sources of carbon. We reasoned that MIT1002-GEM could also provide preliminary insight into how carbon remineralization rates depend upon a heterotroph’s metabolic network, the carbon source used, and environmental conditions. Furthermore, we asked what the model could reveal about the capacity of MIT1002 to recycle nitrogen for the benefit of *Prochlorococcus*.

**Figure 4:**
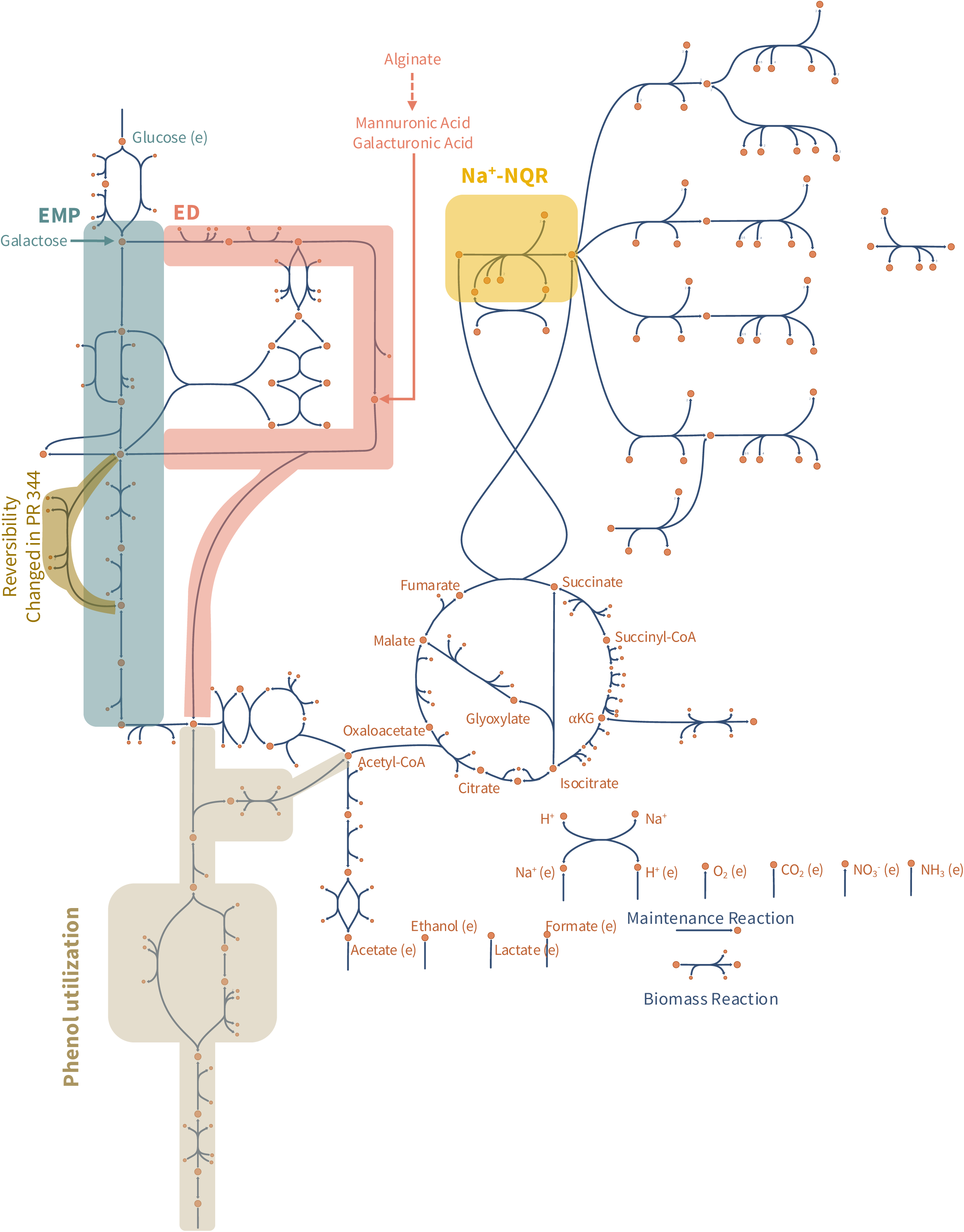
An Escher map of central metabolism and selected pathways of the MIT1002 model. Metabolites in the model are denoted by an orange circle and reactions by blue arrows. The glucose e”change reaction is in the upper left, leading to the two glycolytic pathways: the Embden-Meyerhof-Parnas (EMP) pathway (teal) and the Entner-Doudoroff (ED) pathway (pink). The Pentose-Phosphate Pathway sits between the EMP and ED. A non-phosphorylating glyceraldehyde-3-phosphate dehydrogenase (GAPDHN, tan) is present in lower glycolysis. MIT1002’s unique phenol degradation pathway (light tan) connects extracellular phenol to pyruvate and acetyl - CoA. The TCA cycle is in the center, and directly above the TCA cycle is the branching electron transport chain. The first step in the ETC is a sodium translocating NADH:quinone o”idoreductase (Na^+^-NǪR, yellow). In the bottom right are important e”change reactions for additional metabolites including o”ygen. Below the e”change reactions are arrows depicting the biomass reaction (all metabolites other than biomass removed) and the ATP maintenance reaction.

#### The Entner-Doudoroff Pathway Supports Sugar Acid Metabolism

In line with MIT1002’s genomic content, our model has both a complete Embden-Meyerhof-Parnas (EMP) pathway and a complete Entner-Doudoroff (ED) pathway (Fig. 4). While the presence of these pathways is well supported, it is not clear why seemingly redundant pathways would be present in an organism’s metabolic network. The EMP pathway (producing 2 ATPs per glucose) is considered the canonical glycolytic pathway due to its ubiquity among eukaryotes. However, prokaryotes have several alternative glycolytic pathways, with the ED pathway (producing 1 ATP per glucose) being the most common (83). A possible reason is that, due to its higher thermodynamic favorability, the ED pathway requires less enzymatic protein to catalyze the same flux, allowing cells to invest more protein in biosynthetic pathways driving growth (83). This tradeoff cannot be captured in traditional FBA simulations, as FBA does not include proteome constraints, therefore the higher amount of ATP produced by the EMP pathway directly increases the growth rate and is seen as optimal by FBA (Fig. 5A).

**Figure 5:**
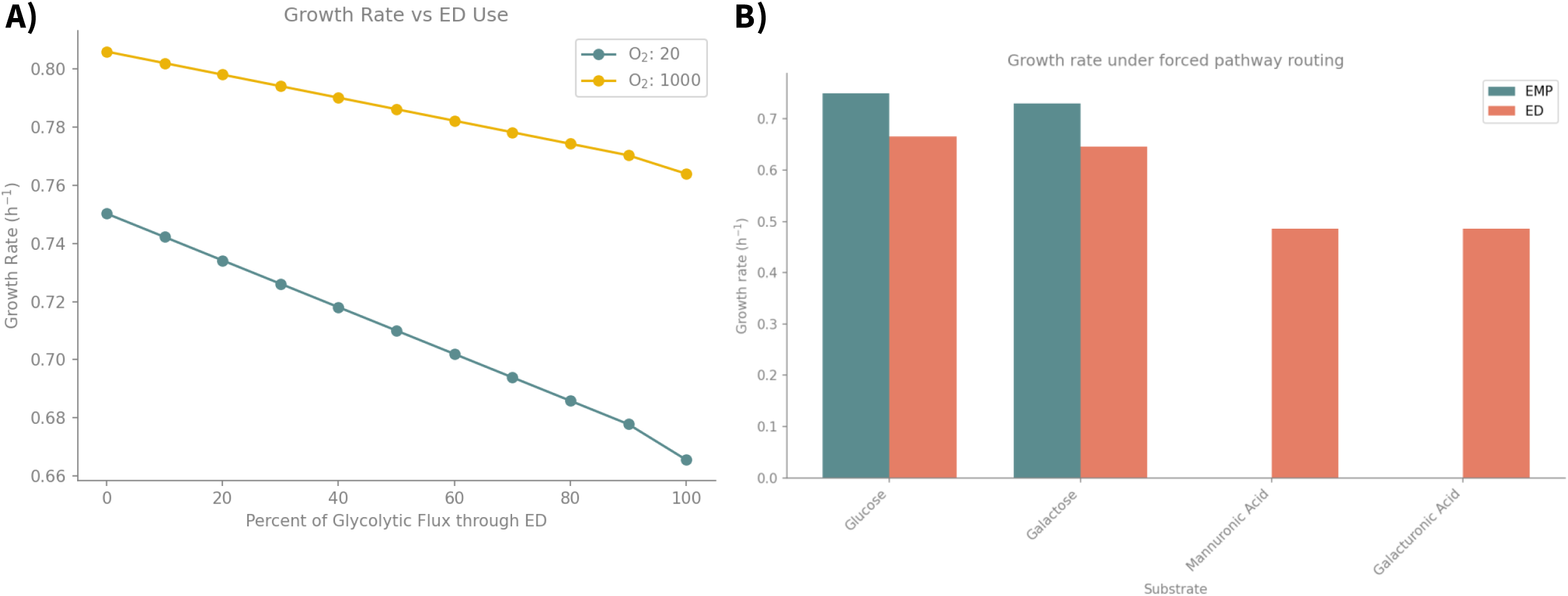
EMP vs ED utilization. A) FBA predicted growth rate for growth on glucose (maximum uptake of 10 mmol glucose / gDW / h) as flux was forced through the ED pathway (by setting a positive lower bound on KDPG-aldolase, rxn03884_c0), under two different oxygen conditions: maximum uptake of 20 mmol O2 / gDW / h (teal), or 1000 mmol O2 / gDW / h (yellow). As flux was forced through the ED pathway growth rate steadily declined for both oxygen conditions with a sharper drop off above 90% of flux forced through ED. B) FBA predicted growth rates on a variety of carbon sources with glycolytic flux forced through the EMP pathway (teal, by setting a positive lower bound on g6p-isomerase, rxn00558_c0) or the ED pathway (pink, as described above).

Alternatively, others have hypothesized that the ED pathway allows the breakdown of sugar acids that cannot be metabolized in the EMP pathway (84). We used MIT1002-GEM to simulate growth on different carbon substrates, forcing all glycolytic flux through either the EMP or the ED pathway (Fig. 5B). When simulating growth on glucose and galactose, the EMP route yields higher growth rates. However, for growth on mannuronic acid and galacturonic acid, the ED pathway is the only option, as forcing flux through the EMP pathway does not support growth, in line with this alternative hypothesis (Fig. 5B). These carbon sources are relevant for MIT1002 as mannuronic acid is one of the monomeric building blocks of alginate (13, 29, 85, 86), and galacturonic acid is an intermediate of pectin degradation (13, 87–89), both of which are nutrient sources in the ocean (90, 91).

#### Carbon Source Degree of Reduction Correlates with Carbon Use Efficiency

Heterotrophic metabolism is a significant factor in the marine carbon cycle because heterotrophic bacteria restructure the DOC pool and regenerate CO_2_. Carbon use efficiency (CUE) quantitatively describes this key role of heterotrophs. To understand how CUE depends on the carbon substrates used, we simulated growth on a wide variety of carbon sources with excess oxygen and used the FBA-predicted fluxes to calculate CUE. In these simulations, all carbon taken up is allocated to either biomass or CO_2_, and so CUE is equivalent to the fraction of carbon taken up that is allocated to biomass. Across these substrates, CUE ranges from 0.41 to 0.72 (Fig. 6A). This broad range of predicted CUE values overlaps with the wide range of efficiency values measured *in situ* in the ocean (from 0.01 to 0.69) (92). Furthermore, this range highlights the potential role of carbon substrates to explain the diversity of efficiency values reported.

**Figure 6:**
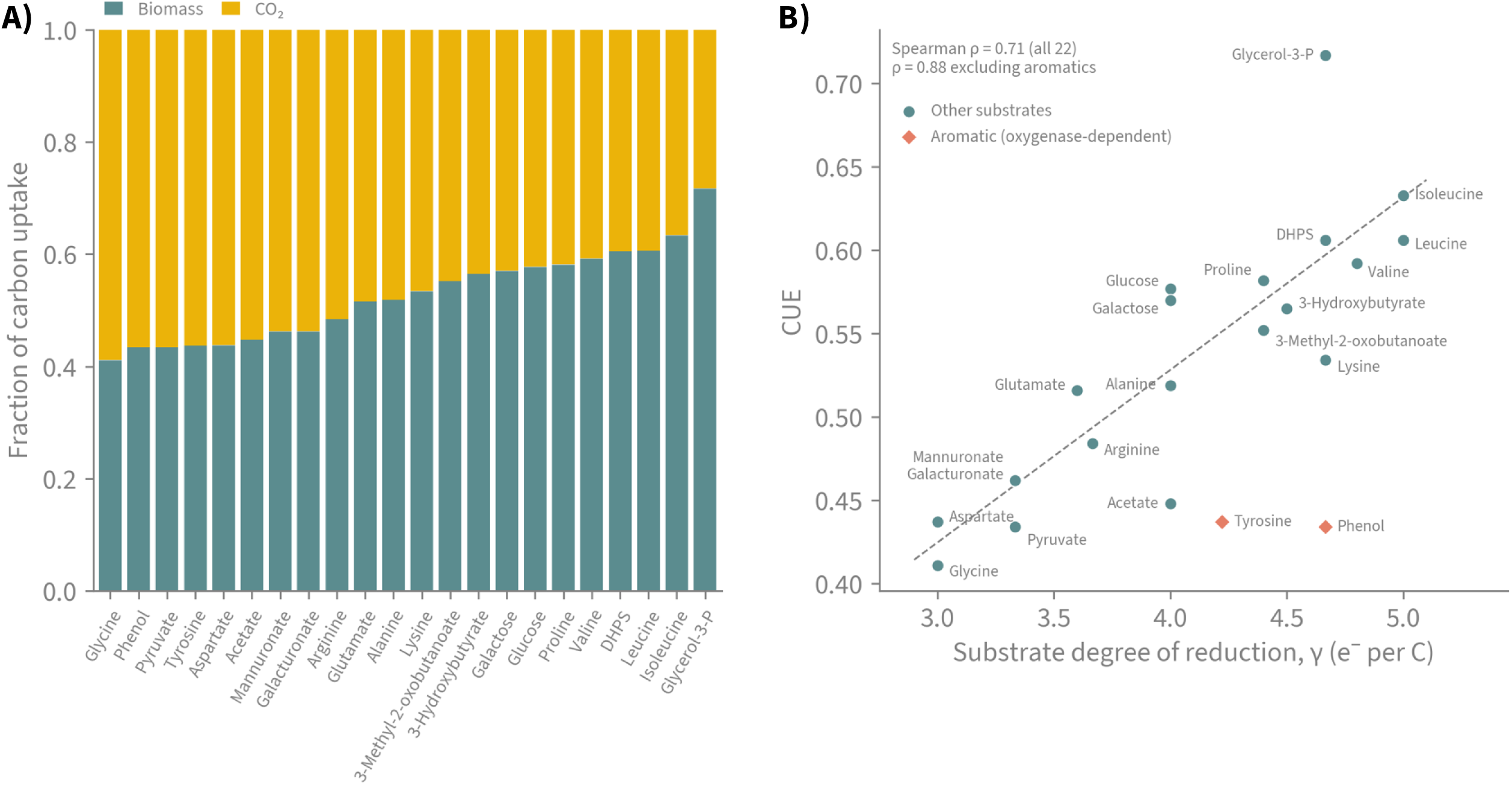
FBA-predicted Carbon Use Efficiency (CUE) for MIT1002 when grown on a variety of carbon sources with non-limiting oxygen (maximum oxygen uptake rate set to 50 mmol/gDW/h). The carbon source uptake rate is set to 60 mmol C/gDW/hr (e.g., glucose is 10, acetate is 30). A) Fluxes of carbon (normalized to the uptake rate) to biomass (teal) and CO2 (yellow=CO_2_). CUE is the height of the teal bar. Bars are ordered by increasing CUE. B) There is a strong positive correlation (Spearman ρ=0.71) between FBA-predicted CUE and the carbon substrate’s degree of reduction (γ). Removing the aromatic substrates tyrosine and phenol from the correlation further strengthens the correlation (Spearman ρ=0.88) as the breaking of the aromatic ring oxygenates the substrates without being used for energy generation.

We found that the CUE was correlated with the carbon substrate’s degree of reduction (the number of available electrons per carbon atom; *γ*) over the range of substrates tested (Spearman ρ=0.71). More oxidized substrates (a lower degree of reduction), such as glycine, result in a lower CUE compared to more reduced substrates (a higher degree of reduction), such as leucine (Fig. 6B). Two notable exceptions to this pattern are the aromatic substrates, phenol and tyrosine, which have a much lower CUE than their *γ* alone would predict. Their lower predicted CUE is due to the oxygenase activity required to break the aromatic ring during catabolism: the electrons pass to O_2_ without going through the electron transport chain (see also Fig. S6). Removing them from the correlation further strengthens the relationship found between CUE and *γ* (Spearman ρ=0.88). The positive relationship between CUE and *γ* over low and intermediate *γ* values is consistent with extensive empirical and theoretical literature (93–95), though previous studies have documented a plateau at higher *γ* values (79, 93, 95, 96).

#### Alteromonas Exudes Ammonia During Growth on Amino Acids

We also used MIT1002-GEM to investigate potential metabolic interactions between MIT1002 and *Prochlorococcus*. We simulated MIT1002’s growth on a variety of carbon sources (Fig. 7), including eight that are among the most abundant exometabolites released by *Prochlorococcus* (97). In simulations of growth on amino acids, FBA predicts that MIT1002 uses amino acids as both a carbon and a nitrogen source, taking up no additional inorganic nitrogen source, despite it being present in the medium (Fig. S7). The model also predicts that growth on amino acids leads to uptake of excess nitrogen, causing the organism to release NH_3_ to the environment. This result corroborates recent experimental evidence that *Alteromonas* remineralizes dissolved organic nitrogen (79). Given that *Prochlorococcus* exports a number of amino acids during growth, it is possible that *Alteromonas* acts as an essential partner by generating a supply of inorganic nitrogen that supports the cyanobacterium’s requirements (19, 98).

**Figure 7:**
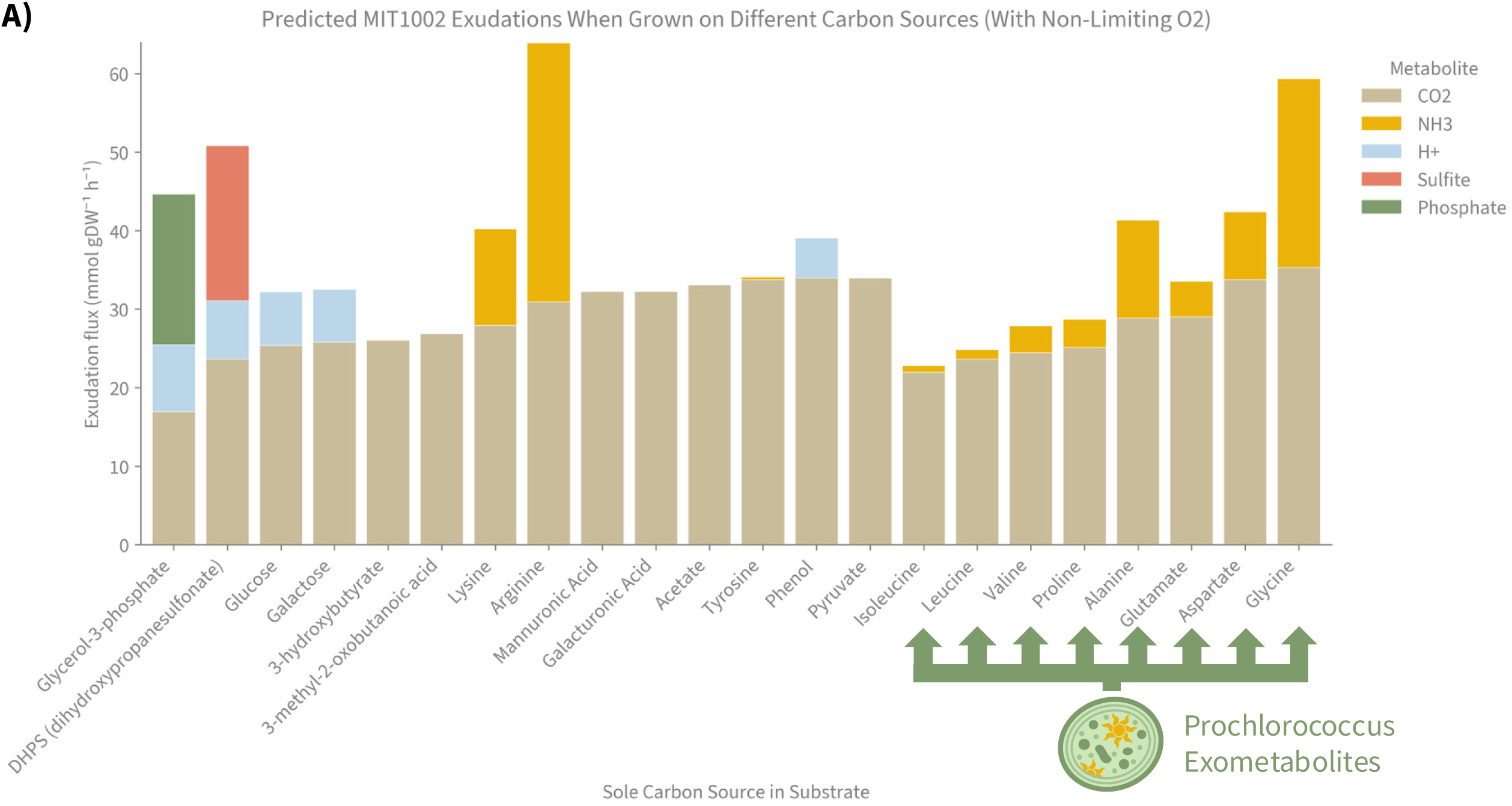
Secretion rates (in mmol/gDW/h) of MIT1002 exudates when grown on different carbon sources. Carbon sources measured in top exometabolites of Prochlorococcus are marked with a green arrow. When growth is simulated on these metabolites, and all other amino acids, MIT1002 secretes NH_3_.

## Discussion

With MIT1002-GEM, we have added a new model to the small set of manually curated metabolic models available for marine heterotrophic bacteria (99–101). In addition to manually curated models such as these, automated reconstruction methods have been used to produce thousands of models for marine heterotrophs, which have yielded insight into the functional guilds of the diverse ocean microbiome (50). These two approaches complement each other in a number of ways. First, high-quality manually curated models, with their embedded experimental data, serve as a ground truth to evaluate the performance of automated reconstruction tools. Second, as automated construction relies on curated templates and reaction databases, manual curation of a single model from that base can reveal errors that are present in all models made from them. For example, we did not find the Na^+^-NQR reaction in the ModelSEED reaction database, or in several other reaction databases (e.g., KEGG), so regardless of its presence in MIT1002’s genome, the initial automated reconstruction could not include it. Because this reaction is thought to be widespread in marine bacteria (80), we expect that other models are missing this reaction. In return, advances in automatic reconstruction are beneficial to manual curation, as manually curated models typically start with draft models made with automated reconstruction tools.

While GEMs are often discussed as tools for simulating growth and predicting reaction fluxes, they are also valuable for generating hypotheses and testing our knowledge about metabolic functions (71, 102). Using MIT1002-GEM we have generated testable hypotheses about the SMF, the ED pathway, cross-feeding with *Prochlorococcus*, and CUE.

Based on genome annotation, we predict that MIT1002 translocates sodium during aerobic respiration to generate a SMF that it can utilize for energy generation, material transport, and motility. The presence and role of the Na^+^-NQR reaction could be tested experimentally by evaluating growth and motility on a range of Na^+^ concentrations or using known NQR inhibitors (e.g., HQNO or korormicin (103, 104)). The presence of Na^+^-NQR also led us to speculate about other potential roles of the SMF, such as genetic regulation. In other marine bacteria, gene expression is coupled with the SMF (e.g., for virulence factors in *V. cholerae* (81)), which may also be the case for MIT1002.

Our results suggest that the ED glycolytic pathway is used for sugar acid metabolism, rather than for breaking down glucose. This hypothesis could be tested by assessing the growth of MIT1002 mutants with gene knockouts in the EMP and ED pathways on sugars and sugar acids or by directly measuring fluxes through reactions in those pathways during growth on different substrates. The alternative hypothesis, that the ED pathway is more favorable for all substrates (including glucose) due to its higher thermodynamic favorability, could be tested with extensions of FBA such as Constrained Allocation FBA (105) or thermodynamics-based FBA (106).

We hypothesize that MIT1002 recycles nitrogen for *Prochlorococcus* by releasing ammonia when using amino acids as a carbon and nitrogen source, as is known to happen more generally between heterotrophs and phototrophs (19, 98, 107, 108). The rate and efficiency of this recycling could be explored computationally using a simulation of a co-culture of the two organisms, e.g., using dynamic FBA (dFBA) with MIT1002-GEM and existing GEMs for *Prochlorococcus* (109, 110), or experimentally using isotopically labeled nitrogen (e.g., ^15^N-labelled amino acid). Notably, other benefits *Prochlorococcus* is known to receive from MIT1002 would require additional modules on top of traditional FBA. For example, to model H_2_O_2_ buildup in the environment and the resulting oxidative stress experienced by *Prochlorococcus* or the cost to MIT1002 to express catalase to detoxify the environment, additional objective functions (111) and proteome constraints (105) could be added to traditional FBA. Similarly, MIT1002’s degradation of proteins by extracellular proteases cannot be easily modeled in standard FBA but could be incorporated in COMETS simulations (112, 113) that take into account proteome constraints and costly production of specific extracellular enzymes (114) or in other coarser-grained models (115).

The wide range of CUE on different carbon sources (Fig. 6A) and the correlation between CUE and γ (Fig. 6B) is expected (38, 92–95). Our results suggest that the relationship between CUE and γ is dependent on details of metabolic pathways. For example, we hypothesize that aromatic substrates, such as phenol and tyrosine, have a lower CUE than expected from the substrate’s γ alone, due to oxygenase activity in their catabolic pathways (Fig. S6). This could be tested experimentally by measuring CUE (116) for MIT1002 when grown on phenol and a non-aromatic substrate with a similar degree of reduction (e.g., lysine or glycerol) and verifying that the CUE when grown on phenol is lower. Furthermore, discrepancies between experimental results and the FBA-predicted CUE values or ordering for a wider range of substrates could reveal non-stoichiometric constraints on CUE (e.g., regulation or proteome allocation). A better understanding of the controls of CUE could be used to better represent heterotrophic bacteria in biogeochemical models, which currently often use a fixed value between 0.15 and 0.6 (117).

The simulations presented here highlight that ecologically significant processes carried out by marine heterotrophic bacteria (e.g., carbon remineralization and interactions with phototrophs) are dependent on details of their metabolic networks. We anticipate that new efforts to better represent heterotrophic metabolism within large scale ocean and Earth-system models will greatly benefit from the availability of high-quality GEMs. In addition to directly incorporating GEMs into larger-scale models (110, 118), GEMs could quantitatively characterize the diversity of microbial heterotrophic lifestyles and distill new rules for improved representation in ecosystem-level models.

## Acknowledgements

We are grateful to members of the Segrè lab for helpful feedback on the manuscript, and to many members of the National Science Foundation (NSF) Center for Chemical Currencies of a Microbial Planet (C-CoMP) for many helpful conversations about *Alteromonas* and GEM curation. We acknowledge support from the NSF C-CoMP Science and Technology Center (OCE-2019589), the NSF–US-Israel Binational Science Foundation (BSF) (grant 2246707), the Human Frontiers Science Program (grants RGP0020/2016 and RGP0060/2021), and the Simons Foundation (grant 542391). H.S. was also supported by the Boston University Kilachand Multicellular Design Program and by National Institutes of Health predoctoral training grant T32GM100842. This is the NSF Center C-CoMP publication #108.

